# Surveillance of H5 HPAI in Michigan using retail milk

**DOI:** 10.1101/2024.07.04.602115

**Authors:** Adam S. Lauring, Emily T. Martin, Marisa Eisenberg, William J. Fitzsimmons, Elizabeth Salzman, Aleda M. Leis, Sarah Lyon-Callo, Natasha Bagdasarian

## Abstract

An outbreak of highly pathogenic avian influenza virus (HPAI) H5N1 in dairy cattle in the United States has spread to at least 136 dairy herds in 11 states. Due to a lack of testing, it has been difficult to determine the number of farms affected and the extent of intrastate spread. We report a pilot study of real time RT-PCR (rRT-PCR) detection of influenza A virus (IAV) in milk purchased from local markets across Michigan. Two out of 13 samples were positive for IAV nucleic acid, suggestive of HPAI H5N1 infection in the originating herds. This small study of a convenience sample suggests that milk-based surveillance might be useful if applied systematically either at dairy processors or point of sale.

## INTRODUCTION

The first documented case of bovine infection was confirmed in March 2024 and as of July 3, 2024, HPAI H5N1 has spread to over 100 herds in 11 States [1]. Michigan received a shipment of infected, asymptomatic dairy cows from Texas on March 8, 2024 [2] and as of July 2024, HPAI H5N1 has spread to 25 dairy herds and eight poultry farms across 11 counties in Michigan [1 and unpublished data].

While the interstate spread of HPAI H5N1 in dairy cattle has been well documented through traditional and genomic epidemiology [3,4], the amount of farm-based testing has made it challenging to define the extent of spread within individual states and the cows and herds affected. Current surveillance is dependent on farm owners reporting and allowing testing of ill animals. Coupled with the current surveillance strategy, a more granular method of surveillance could be critical for assessing the overall trajectory of the outbreak and the effectiveness of mitigation measures on farms.

On April 25, 2024, the United States Food and Drug Administration reported that up to 20% of milk products sampled across the country were positive for influenza A virus (IAV) nucleic acid by real-time reverse transcription polymerase chain reaction (rRT-PCR). Infectious virus has not been detected in milk subjected to standard pasteurization procedures [5]. Since this original announcement, several groups have reported detections of IAV nucleic acid in retail products in the US Midwest [6] and Canada [7]. Here, we describe a pilot program of testing pasteurized milk from local, farm-affiliated retailers for local surveillance of the HPAI H5N1 in the State of Michigan.

## METHODS

### Sample Collection

Milk was purchased from local dairy stores across Michigan (sites indicated in Table 1) and stored at 4°C.

**Table 1.** Origin and rRT-PCR test results for 13 milk samples.

| Sample | County of Dairy | County of Purchase | Plant Code | Purchase Date | SIV Ct | Xeno-Control Ct | Result |
| --- | --- | --- | --- | --- | --- | --- | --- |
| 1 | Monroe | Washtenaw | N.D | 6/8/2024 | Undetermined | 30.85 | Negative |
| 2 | Allegan | Washtenaw | N.D | 6/8/2024 | Undetermined | 29.67 | Negative |
| 3 | Macomb | Macomb | N.D | 6/8/2024 | Undetermined | 30.95 | Negative |
| 4 | Oakland | Oakland | 26-907 | 6/8/2024 | Undetermined | 30.74 | Negative |
| 5 | N.D. | Ingham | 26-306 | 6/16/2024 | 36.65 | 31.02 | Positive |
| 6 | Osceola | Kent | 26-136 | 6/16/2024 | Undetermined | 30.83 | Negative |
| 7 | Osceola | Kent | 26-136 | 6/16/2024 | Undetermined | 30.85 | Negative |
| 8 | N.D. | Ingham | 26-306 | 6/7/2024 | 37.55 | 30.44 | Positive |
| 9 | Allegan | Allegan | N.D | 6/7/2024 | Undetermined | 30.16 | Negative |
| 10 | N.D. | Ingham | 26-111 | 6/7/2024 | Undetermined | 29.98 | Negative |
| 11 | N.D | Ingham | 26-111 | 6/7/2024 | Undetermined | 29.95 | Negative |
| 12 | Oscada | Washtenaw | 26-034 |  | Undetermined | 30.59 | Negative |
| 13 | Marquette | Alger | 26-228 | 6/16/24 | Undetermined | 30.65 | Negative |

N.D. = no data
Highlighted cells indicate county with impacted dairies (as in text)

### RNA harvest and rRT-PCR

RNA was extracted from 200µl of each milk sample using the MagMAX Viral/Pathogen II (MVP II) Nucleic Acid Isolation Kit (ThermoFisher, A48383) on a Kingfisher Machine; RNA was eluted in 50µl of elution buffer. Duplicate mock-purified samples were harvested using 200 µL 1x PBS. Two microliters of undiluted Xeno RNA extraction control were added per sample. The VetMAX-Gold SIV Detection kit (ThermoFisher, 4415200) was used for real time reverse transcription polymerase chain reaction (rRT-PCR) with 8 µL of input RNA. Duplicate positive controls were run using 8 µL of Influenza Virus-Xeno RNA Control Mix and duplicate “no template” controls were run using 8 µL of nuclease-free water. Run reporter dye thresholds were set at 5% of the average maximum fluorescence value of the amplification signal of the positive control reactions at cycle 40.

## RESULTS AND DISCUSSION

We used a “convenience sampling” strategy in which individuals from laboratories at the University of Michigan purchased milk from locations geographically distributed across the State of Michigan during personal trips. Local dairy stores were specifically targeted, as opposed to large supermarket chains, in order to sample milk processed near the farm of origin. The purchase date, dairy county, county in which the milk was purchased, and the plant (i.e., processor) code for the 13 samples are shown below (Table 1).

We extracted RNA from 200 µl of each sample and detected influenza A virus nucleic acid using a Swine Influenza Virus RNA Test Kit. The primers and probes in this kit are broadly reactive against animal-origin IAV, and a positive signal for IAV in cow’s milk was interpreted as strongly suggestive of HPAI H5N1 infection in one or more of the source farms. The cycle threshold value (Ct) for the Xeno RNA extraction control was in the expected range (Ct 27.5-34, per manufacturer) as were the Ct values for the positive (27.42; expected Ct 25-29 per manufacturer) and negative control (undetermined) samples (Table 1). Two out of the 13 samples were positive with Ct values of 36.65 and 37.55, respectively. While purchased on different days, both had the same processing plant code and appeared to be from Ingham County. Testing for infectious virus was not performed. All sampled milk products had been pasteurized, which has been shown to effectively inactivate HPAI H5N1 viruses.

Counties with impacted dairies as of July 2, 2024, include Montcalm (2), Ionia (5), Isabella (2), Ottawa (2), Barry (1), Gratiot (5), Ingham (1), Allegan (1), Clinton (5), Calhoun (1)

In this pilot study, we were able to detect presumptive H5N1-derived nucleic acid in two milk samples derived from the same processing facility, both with links to a county with known positive dairy herds. Two other samples from the same county were negative, and all milk samples from counties without documented positive herds, tested negative.

This suggests that more systematic surveillance of milk from processors or individual dairies could be used to determine the extent of within-state spread over time. Broadly implemented, this would fill a critical surveillance and provide key data for assessing the effectiveness of mitigation measures.

## ACKNOWLEDGMENTS

We thank Richard Webby and Andrew Bowman for suggestions on extraction of RNA from milk and RT-PCR protocols. Funded by discretionary funds available to Dr. Lauring.

